# Onsager’s variational principle in proliferating biological tissues, in presence of activity and anisotropy

**DOI:** 10.1101/2023.09.01.555879

**Authors:** Joseph Ackermann, Martine Ben Amar

## Abstract

A hallmark of biological cells is their ability to proliferate and of tissues their ability to grow. This is common in morphogenesis and embryogenesis but also in pathological conditions such as tumour growth. To consider these tissues from a physical point of view, it is necessary to derive fundamental relationships, in particular for velocities and density components, taking into account growth terms, chemical factors and the symmetry of cells and tissues. The aim is then to develop a consistent coarse-grained approach to these complex systems, which exhibit proliferation, disorder, anisotropy and activity at small scales. To this end, Onsager’s variational principle allows the systematic derivation of flux-force relations in systems out of equilibrium and the principle of the extremum of dissipation, first formulated by Rayleigh and revisited by Onsager, finally leads to a consistent formulation for a continuous approach in terms of a coupled set of partial differential equations. Considering the growth and death rates as fluxes, as well as the chemical reactions driving the cellular activities, we derive the momentum equations based on a leading order physical expansion. Furthermore, we illustrate the different interactions for systems with nematic or polar order at small scales, and numerically solve the resulting system of partial differential equations in relevant biophysical growth examples. To conclude, we show that Onsager’s variational principle is useful for systematically exploring the different scenarios in proliferating systems, and how morphogenesis depends on these interactions.

## 1 Introduction

Growth of biological materials can occur *in vivo* or in *in vitro*. Proliferative cells are mainly responsible for the growth process. *In vitro* experiments mainly involve epithelia on a substrate, cysts or aggregates in setups with nutrients. Only certain cells form such colonies, mostly those with a cancerous or stem character. When the number of cells becomes very large, discrete models, such as the vertex models, are limited by the size of the simulations and a continuous approach or coarse-grained approach may be more appropriate [1]. The relation between the microscopic description of passive and active tissues and their hydrodynamic equations have notably been more understood in the last years [2, 3]. For *in vivo* growth processes, we face the same limitations but in addition there are more partners of different types such as cells, tissues and fibres. The biological properties of the system depend on more complex interactions between the different partners, and the interactions between them are not always well established and quantified. The aim of this paper is therefore to develop a consistent formalism for the evolution of a proliferating population of cells undergoing chemical reactions and activities. These cells may have specific physical properties at small scales, such as a nematic or a polar order, and our formalism will incorporate such properties at the level of a continuous approach.

Three examples will be chosen to make this formalism more concrete: the growth of cancerous aggregates, growth of epithelia *in vitro*, and growth of interconnected tissues. The last case more closely mimics the tumour growth in its environment *in vivo*.

Indeed, the key point of our approach is the coupling of growth with activity or, more generally, with chemical reactions. In fact, growth requires the consumption of energy as well as nutrients. This energy is produced by aerobic respiration and is necessary for the cells to carry out their functions. In proliferating cells, such as stem cells or cancer cells, cell energy is necessary for cell division [4]. Because of their abnormal proliferation, cancer cells are the best example of large energy consumers. The process of energy production can be divided into two steps: first, the production of ATP molecules from ADP and storage (this step is related to respiration), second the hydrolysis of ATP and production of mechanical energy [5]. At the same time, proliferation requires nutrients (such as oxygen, glucose…), so the nutrient consumption and the proliferation rates are coupled to each other. Polarity and nematic properties are also important players in cell proliferation, as they are often crucial for understanding single cell or collective cell behaviours [6, 7]. Cells in epithelia are highly polarised, in particular due to an asymmetry in cell shape, that gives them their apical-basal polarity. In epithelia, cells show an asymmetry at the origin of their apical-basal polarity. We show here that this polarity, observed at the scale of a single cell, determines the shape of a multicellular epithelium. A typical example concerns the cysts of stem cells in the human epiblast phase in fetal life or in experiments *in vitro* with human pluripotent stem cells [8] as shown in Fig. 1(a). In contrast, cancer cells of epithelial origin lose their polarity and soften their cytoskeleton (the ensemble of the cell filaments) during the EMT transition. This process promotes metastasis by facilitating their escape from the epithelium. However, a nematic order may exist in the tumour microenvironment due to nematic cells such as fibroblasts or tissue components such as collagen fibers, see Fig. 1(c). A tumour and its stroma represent a highly dynamic mixture of active species, and cancer cells exhibit a wide variety of behaviours, leading to different spatial structures in the tissue or organ host. Different long-term evolutions are expected, which need to be represented by a coarse-grained approach.

**Figure 1.**
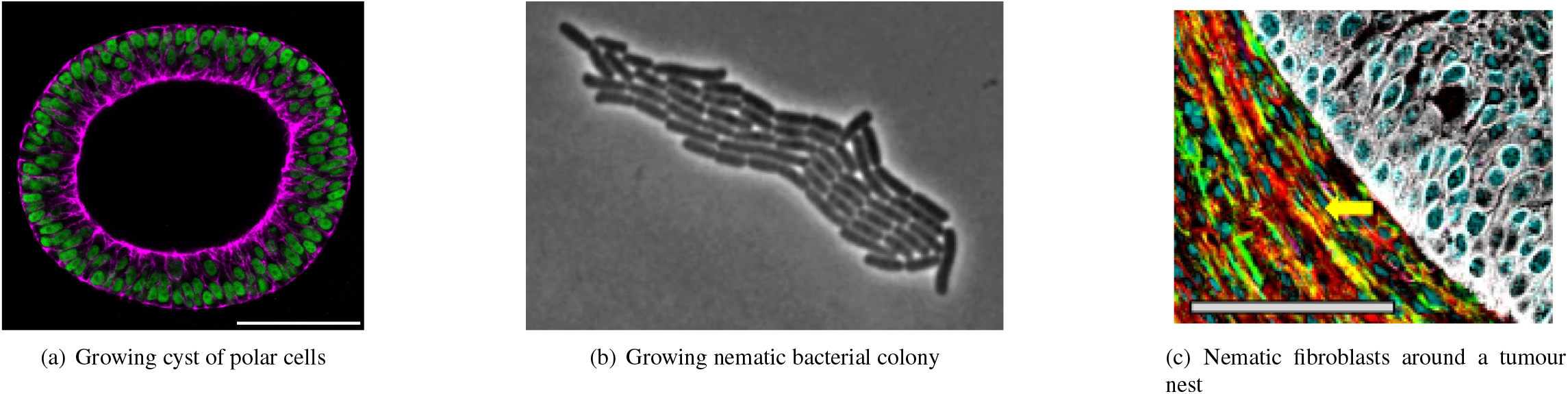
**1(a)**: Growing cyst of polar cells (from [8]). Polarity induces an anisotropy in growth, which in turn creates stresses that have a feedback effect on the growth. **1(b)**: Growing bacterial colony (from [11]). The nematic nature of the bacteria leads to a specific structure in the colony, and in the polar field. **1(c)**: Cancer cells may not have an intrinsic polar or nematic order. However, in desmoplastic tumours, as shown the environment is highly ordered, as shown here with fibroblasts surrounding a lung tumour nest (from [12]).

**Figure 2.**
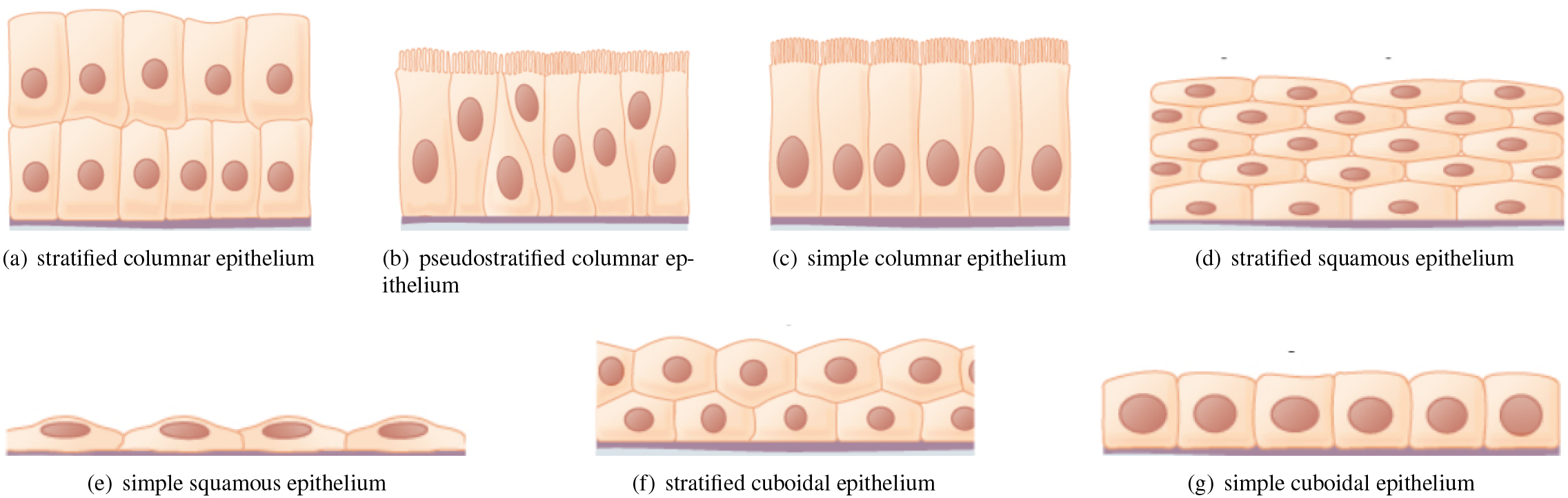
Different epithelia depending of the shape of cells (images from [33]). Epithelia can be stratified, pseudo-stratified or simple. Cells can have a clear polarity in their shape and structure (as in ciliated columnar cells), or can be nematic (like the cuboidal cells). In any case, the presence of a basal membrane creates a polarity in the epithelium.

In summary, different cell types coexist in tissues with the so-called extracellular matrix. This requires the integration of multiple components into the model when studying living tissues.

The aim of this paper is to present a formalism that takes into account the different possible phenomena that can occur in a living system. We want to establish the dynamical equations for the mass densities of the different components and their velocities, which are obviously coupled to the growth and to the chemicals that cause the phenotypic changes of the cells. We will also show that the macroscopic patterns strongly depend on the symmetry that the cells exhibit at small scales as polar or nematic order. Using the software Comsol Multiphysics [9], we apply our equations to 2 dimensional systems, as most experimental efforts have focused on these systems, as a first step towards more complex 3D modelling [10].

The article is organised as follows: First, in section 2, we present Onsager’s variational principle. In section 3, it is applied to apolar and non-nematic proliferating cells and activity is further added in section 4. In the following sections 5-6, the growth of nematic cells is treated with this formalism and in section 7 it is applied to polar cells. Finally, examples of growth mixtures with three components and in the presence of activity and nematic order are shown in section 8.

## 2 Onsager’s variational principle

Onsager’s variational principle has already been applied in soft matter [13, 14, 15], and in active matter to describe tissues [16, 17, 18], in particular cancerous tissues [19, 1]. Here, we briefly review the methodology of the Rayleigh-Onsager’s variational principle [20, 21]. First we write a free energy function *ℱ* of the state variables. Then we identify the fluxes in the conservation equations and we construct a quadratic function in these fluxes, called the dissipation function. Finally, the sum of the free energy variation and the dissipation, defined as the Rayleighian *ℛ*, is minimised with respect to the fluxes *X*, yielding flux-force relations:

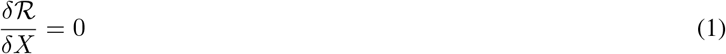

These flux-force relationships arise from the fact that the free energy can be written as quadratic in the fluxes and in the state variables close to the equilibrium [21]. Further from equilibrium, the relaxation of the flux-force relations can be written as:

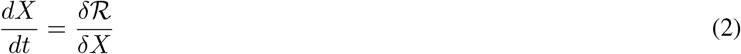

The momentum equation can therefore be written as follows:

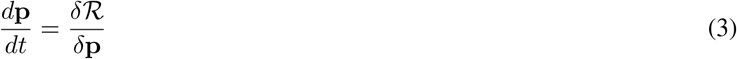

where **p** is the momentum of the center of mass. The main idea of this article is the fact that the proliferation rate is a flux, so it can be derived by extremising the Rayleighian, and it can be coupled to other fluxes [20]. Moreover, since growth is highly dependent on the structure of the tissue, it is indirectly governed by the various couplings that give rise to the architecture of the tissue, even in the absence of direct coupling between the growth and the other fluxes in the flux-force relations.

## 3 Application to non polar cell mixing in a fluid

### 3.1 Fundamental evolution equations

We first describe the concept of homeostasis which states that cells adjust their density to the optimal density in terms of their free-energy. We consider only a mixture of cells of mass fraction *ϕ* and its environment of mass fraction 1*−ϕ*. This free energy *ℱ* includes for example attraction/repulsion interactions between cells and volume exclusion. We also introduce the cell chemical potential *µ*. As a first step, we do not introduce any chemicals into the system, which behave as if cells were passive species. We write the conservation equation for the cell mass fraction *ϕ* in the mixture, with a local velocity **v** and a growth rate *G*. The fluxes in the system are *G* and **v**, and a dissipation function *W* is written in terms of the quadratic couplings between fluxes. The friction between species is calibrated by a friction coefficient *ξ* and the dissipation in the cell phase is measured by the viscosity *η*. A growth coefficient *g*_0_ will be related to the time scale of the growth. A coupling between growth and velocity is also introduced and controlled by a parameter *g*_1_, and it turns out that the growth rate can be interpreted as a pressure through this *g*_1_ parameter. In this presentation, we limit ourselves to a single cell type, the extension to multiple cell types is straightforward and postponed at the end in section 8. The different equations read:

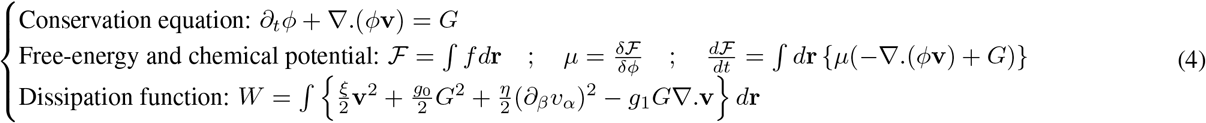

Defining 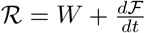 and minimizing *ℛ* with respect to the fluxes, it comes:

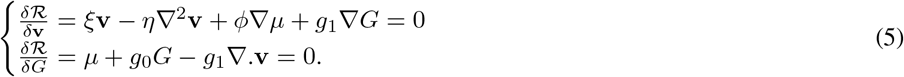

From Eq. 5, we get *G* and a momentum equation:

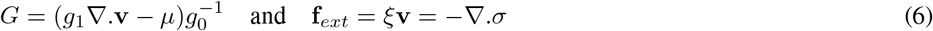

where the elastic stress is:

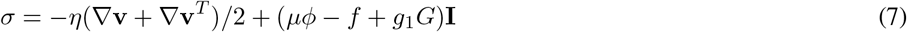

**I** being the unit tensor. The second equation on right of Eq. 6 is equivalent to a Darcy law. Two terms contribute to *G* in Eq.6: the divergence of the velocity and the chemical potential. Consider a tissue which occupies the semi-negative plane with a cell flux of increasing velocity towards the boundary: then, the cell growth rate *G* will increase. Conversely, if the velocity decreases in the direction *e*_*x*_, then cells die. For the second contribution in *G*, we need to define the free-energy density *f*.

### 3.2 The Cahn-Hilliard free-energy density

We choose for *f* a Cahn-Hilliard free-energy density, which includes a Flory-Huggins free-energy for the homogeneous part and a square gradient energy in *ϵ*^2^, that represents an interfacial free-energy. This energy includes the diffusion of the species in the mixture, as well as the volume exclusion, the attraction/repulsion between cells, an energy cost linear in the mass fraction, and the cost for sharp spatial variation of cell volume fraction. The free-energy density writes:

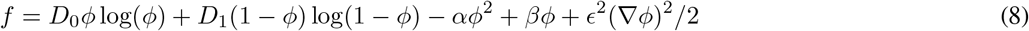

and we deduce the cell chemical potential:

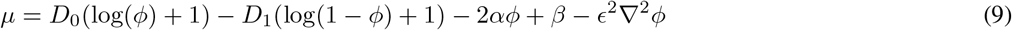

*D*_0_ and *D*_1_ control the cell diffusion and the volume exclusion, *α* controls the attraction/repulsion between cells, and *ϵ >* 0 controls the size of interfaces. The existence of a compact tissue at equilibrium with its environment implies *α >* 0 as cells adhere to each other. Note that, the term proportional to *β* is chosen to be linear, which plays an important role in homeostasis since the cell growth rate *G* depends linearly on *µ* (Eq. 6). At equilibrium, *v∼* 0 and the system tends to stabilise around *G* = *− µ/g*_0_ = 0. Therefore, homeostasis corresponds to an extremum of the free-energy *µ* = 0 for a mass fraction *ϕ* = *ϕ*^*\**^. In the case of a dense phase Ω_1_ with a mass fraction *ϕ*_1_ in equilibrium with a dilute phase Ω_2_ with a mass fraction *ϕ*_2_, the system is in equilibrium if the equalities of the chemical potentials and the pressures are verified : *µ*_1_ = *µ*_2_, Π_1_ = Π_2_. Therefore, if *ϕ*^*\**^ *> ϕ*_1_ the tissue Ω_1_ will invade its surroundings unless the growth rate is negative in the dilute phase or the interface. On the opposite, if *ϕ*^*\**^ *< ϕ*_1_, the tissue will collapse unless the growth rate is positive in the dilute phase or the interface. In the case of *ϕ* = *ϕ*_1_ = *ϕ*^*\**^, the evolution of the growth will depend on the behaviour in the dilute phase and the interface.

Interestingly, growth seems to change the tissue properties in some cases, so that their surface tension is modified [22]. This could be due to an active process involving stresses between proliferating cells and their surrounding. Alternatively, the parameters of the free-energy might depend on the growth rate.

## 4 Non polar cells mixing with activity

We now consider chemical reactions with rate *r* and concentration *c*_*i*_. In the case where the chemical reaction is related to ATP degradation, *r* can be treated as an active term. In this section, although cells are not considered as nematic entities, the system may have an orientation locally characterised by a traceless tensor corresponding to the surface orientation which undergoes an active stress: **Φ**_*αβ*_ defined by **Φ**_*αβ*_ = (*∂*_*α*_*ϕ*)(*∂*_*β*_*ϕ*) *−δ*_*αβ*_(*∂*_*κ*_*ϕ*)^2^*/d*, where *d* is the dimension of the system. In addition, there is also a deviatoric strain rate: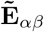. Taking into account this new complexity, Eq. 4 needs to be updated. Note that once we introduce the tensor **Φ** a new coupling appears that is not related to chemicals. We introduce the concentration *c*_*i*_ of molecules, their velocity **v**_*i*_, and their stochiometric coefficient *?*_*i*_ and we define preliminary quantities:

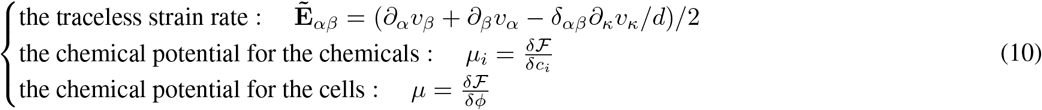

The conservation equations, the free-energy and the dissipation write:

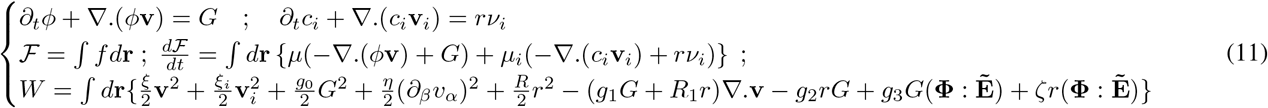

where **A** : **B** = **A**_*αβ*_**B**_*αβ*_ is the tensorial product. Minimising the Rayleighian with respect to the fluxes leads to:

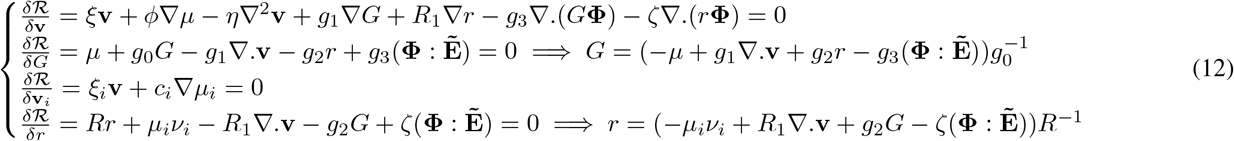

where we have neglected the coupling between the velocities of the chemicals and the other fluxes. In general, the chemical potential for the chemical species can be written as :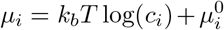, which leads to the Gibbs relation 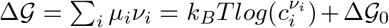 (where is the Gibbs free-energy), and to a diffusion velocity **v**_*i*_ = − (*k*_*B*_*T/ξ*_*i*_)∇ *c*_*i*_. Since proliferation requires ATP production, and that ATP production is stimulated by proliferation, there is a positive coupling exists between them with *g*_2_ *>* 0. It should be noted that the general presence of cell proliferation may inhibit other functions of the cell due to the metabolic burden.

In the following we examine two cases for the growth process. It is controlled either by density or chemical concentrations leading to *G∼ ϕ*(*ϕ*_0_ *−ϕ*) or *G∼ ϕc*_*i*_. These relations come from the dependence of the variable *g*_2_ on the different concentrations and mass fraction and the assumption *g*_1_ = *g*_3_ = 0. Together with this, we assume in those cases that the reaction rate *r* is constant and fixed by the Gibbs term Δ *𝒢*_0_.

In the next section, we investigate the growth of nematic cells.

## 5 Growth of nematic cells

We now discuss growth in the context of active nematic order. Among other examples, we can mention the case of proliferating cancer-activated fibroblasts [23, 12], spindle cell sarcomas [24, 25] or MDCK cells [10], but also the final steps of wound healing performed by myo-fibroblasts [26, 27] or fibrosis occurring in pathologies and around implants [28, 29, 30]. Tissue growth often follows tissue anisotropy, and stresses resulting from this anisotropic growth can have a feedback effect on the growth itself [8]. Therefore, anisotropic growth must be taken into account when modelling cancer growth [31]. Note that the nematic order is a significant player in many biological processes and has been introduced, for example, in the modelling of wound healing and the dynamics of confined colonies of nematic cells [18]. We write the growth equation for *ϕ, c*, and the nematic order matrix **Q**. The nematic matrix **Q** is a locally symmetric and traceless matrix built up from the orientations of the cells. If each cell has an orientation **n**, the orientation matrix writes **Q**_*αβ*_ =*< n*_*α*_*n*_*β*_ *− n*_*γ*_*n*_*γ*_*/d >* where *<>* is an ensemble average and *d* is the space dimension. **Q** is thus an indicator of the orientation order and disorder.

The concentration *c* can either be related to respiration, where *c* is the *O*_2_ concentration, or to ATP degradation. The fluxes are now **v, v**_*c*_, *G*, **Γ**, *r*_*c*_. For simplicity, we do not consider all the possible couplings but select some of them. We consider that the friction is controlled by *ξ*, the viscosity by *η*, the growth by *g*_0_, the shear by the shear parameter *λ*. Two active couplings between the nematic source term **Γ**, the proliferation term *G* and the deviatoric strain rate 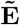 are governed by *ζ*_0_ and *ζ*_1_ (Eq. 15).

We start by defining the free energy, which includes the Cahn-Hilliard Flory-Huggins, the nematic order, and the chemical free energy : *f* = *f*_*F H*_ + *f*_*N*_ + *F*_*c*_. The Cahn-Hilliard Flory-Huggins free energy is written as above according to Eq. 8. The nematic order free energy *f*_*N*_ and the chemical free energy *f*_*c*_ are written as follows:

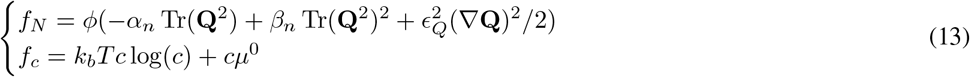

If the orientation of the cells is constrained by boundaries, i.e. the cells spontaneously orient themselves normal or perpendicular to the boundaries, then *f*_*N*_ is coupled to **Φ**_*αβ*_ = (*∂*_*α*_*ϕ*)(*∂*_*β*_*ϕ*) *− δ*_*αβ*_(*∂*_*κ*_*ϕ*)^2^*/d*, already introduced above. In this case the nematic free energy *f*_*N*_ is modified and becomes:

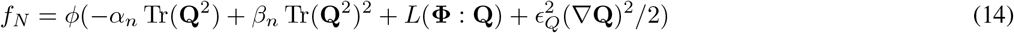

We now explicit the system of dynamical equations for our mixture of nematic cells in the presence of nutrients and reactive chemicals.

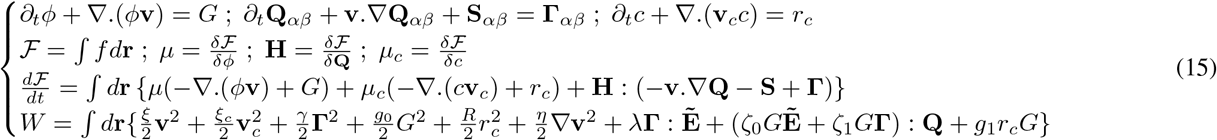

where **S**_*αβ*_ = **Q**_*αk*_**Ω**_*kβ*_ *−* **Ω**_*αk*_**Q**_*kβ*_ and **Ω**_*αβ*_ = (*∂*_*β*_*v*_*α*_*− ∂*_*α*_*v*_*β*_)*/*2, and 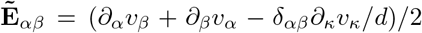. Note that other couplings are possible, such as active coupling of higher order in **Q**, such as *∼* **v**(**Q**.(*∇* .**Q**)), that can have drastic consequences on the stability of the system [32]. In the following, for simplicity, the reaction rate for ATP is considered to be constant and fixed by the chemical potential *µ*_*c*_, with *g*_1_*/R →* 0, and 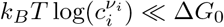, so that *r*_*c*_ *∼ −*Δ*G*_0_.

It comes:

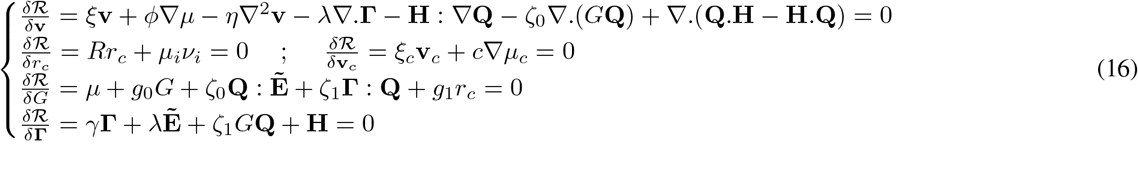

We now apply this formalism to a compact tissue in its surrounding environment in 2 dimensions. This tissue can involve, for example, a tumour nest and the micro-environment which is called the stroma. We study the scenarios for different values of the parameters already introduced.

## 6 Predicted patterns of tissue growth with nematic ordering

Starting from an initial disc of adherent cells, we describe different evolutions with respect to different combinations of energies and dissipation functions. We consider three cases according to the coupling parameters in relation to the simulations reported in Fig. 3. In the following, the tissue is assumed to grow according to a logistic growth function *G ∼ ϕ*(*ϕ*_0_ *− ϕ*) (this implies *ζ*_0_*/g*_0_, *ζ*_1_*/g*_0_ *→* 0, and *−*(*µ* + *g*_1_*r*_*c*_)*/g*_0_ *∼ ϕ*(*ϕ*_0_ *− ϕ*)) (see end of section 4). All of the parameters used in the simulations are shown in Table.Sup. 1.

**Figure 3.**
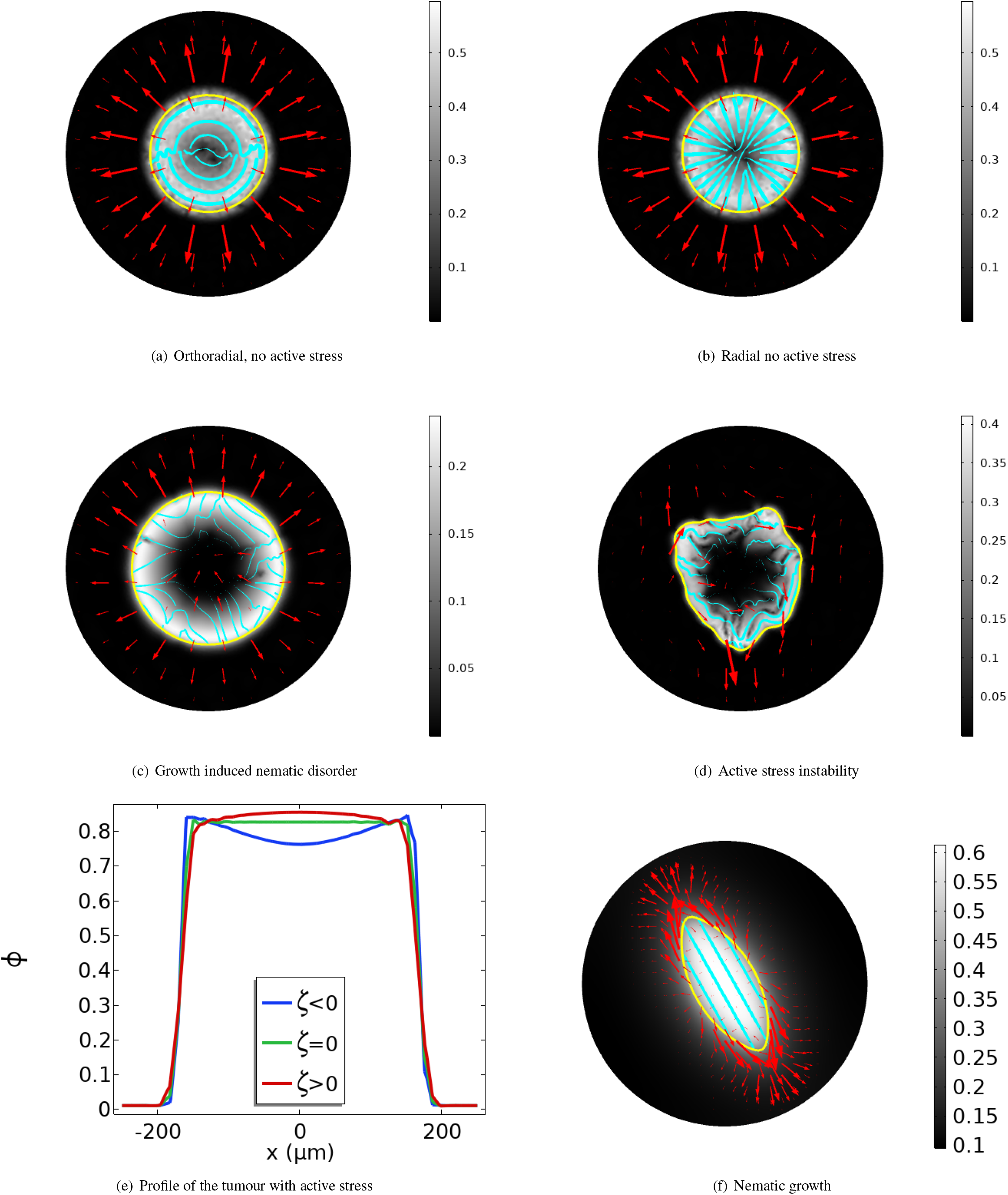
The yellow line corresponds to the boundary of the tumour *ϕ* = 0.35. The red arrows correspond the velocity field. The blue contours correspond to the nematic order. The white-black spectrum correspond to the measure of the nematic order parameter. 3(a): Antiflow-alignement case. 3(b): Flow-alignement case. 3(c): Active and growth induced nematic disorder. No active stress is introduced. 3(d): Growth induced nematic order with active stress. 3(e): Profile of the tumour with active stress. 3(f): Natural nematic aggregate with active growth stress.

### 6.1 Tissue with flow alignment induced nematic order

The orientation of the cells is sensitive to the presence of a flow such as the interstitial flow [34, 35, 36]. It is therefore of interest to study the orientation of proliferating cells under the flow induced by their own proliferation.

We consider the flow alignment introduced by the parameter *λ* = *λ*_0_*ϕ* ≠ 0 in the first equation of Eq. 16. We discard the couplings between the growth and the velocity (represented by 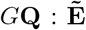) and between the growth and the nematic source term (*G***Γ** : **Q**) in the same system of equations so we assume that *ζ*_0_ = *ζ*_1_ = 0. Nevertheless, a nematic order appears along the velocity field in the growing region and it is stronger at the periphery compared to the center of the tumour. Two cases are shown in Fig. 3 corresponding to *λ <* 0 and *λ >* 0. In the case of an anti-alignment (*λ >* 0), the cells are oriented perpendicular to the velocity field, see (Fig. 3(a)). In the case of flow alignment *λ <* 0, a nematic order appears, i.e. along the velocity field (Fig. 3(b)).

### 6.2 No flow aligment but coupling between the growth and the nematic ordering

We show here that proliferating cells may align along some axis related to the division axis, and stresses that arise from this order can induce an instability. For proliferating or dying cells with no natural orientation, the theory predicts the possibility of spontaneous nematic ordering through the parameter *ζ*_1_, which couples the nematic source term with growth. We therefore focus on the combination of parameters where *ζ*_1_ ≠ 0 but *λ* = *ζ*_0_ = 0. For *ζ*_1_ *<* 0, a nematic order with proliferation spontaneously occurs for *G >* 0 (Fig. 3(c)), and similarly for *ζ*_1_ *>* 0 and *G <* 0. Contrary to the case of flow alignment, the nematic order is only short range because the orientation varies quite strongly in the disc. In fact, in these, cases, we do not observe along-range order over the whole nest in these cases. In addition, the presence of an active stress *ζ*_0_ ≠ 0 leads to instabilities (Fig. 3(d)).

### 6.3 Coupling growth with velocity in for nematic cells

Many types of cells have a spontaneous nematic properties, or can acquire an orientation close to boundaries. Here we study their growth in presence of active stress. We discard the flow alignment *λ* = 0, as well as the coupling between the growth and the nematic source term *ζ*_1_ = 0. However, we introduce a coupling between the growth and the fluid velocity indicated by *ζ*_0_: *ζ*_0_ ≠ 0. We assume that the cells have a natural order close to the boundaries *α*_*n*_ = 0, *L* ≠ 0. This combination of a stress related to growth in the presence of cells spontaneously oriented by the boundaries of the tumour leads to a non-uniform profile of cell density in the tumour (Fig. 3(e)). Indeed, it has been observed that cells can be more concentrated in the middle of the spheroid than at its periphery [37], although it is not always the case (see Ref. [38] in Fig. 1(b) at *t* = 0).

In the case of a natural nematic order (*α*_*n*_ = *α*_*n*0_*ϕ >* 0) and in the presence of a growth-induced stress *ζ*_0_ *<* 0, the growth follows the nematic order, leading to an anisotropic growth, which is well known in bioelasticity (Fig. 3(f)) [39].

## 7 Growth of polar cells

This section examines the growth of polar cells, i.e. asymmetric cells with a clear preferred axis of orientation. Apart from growth, polarity is highly related to motility, and in those cases can be related to actin flows and chemical gradients [40]. Hence interactions between growth and motion can be of interest. We derive the hydrodynamic equations [41] and and focus on the generation of a front instability. This analysis is particularly relevant to epithelial cells, which have the shape of a cuboid with a polarised cytoskeleton and two distinct opposite edges. Pluripotent stem cells are also involved but in this case, they look like a half-cuboid truncated along a diagonal plane, and they organise their proliferation by joining in pair, head to tail, in monolayer cysts in a free space. In this case the epithelium is called pseudostratified. Their growth has been shown to be highly anisotropic [8]. Since most human cancers originate from epithelial tissues, it is crucial to pay attention to this growth category.

Oriented cell division plays an important role in morphogenesis and organogenesis [42]. In epithelia, the plane of division can either be orthogonal to the polarity of the cells and promote a stratification of the epithelium, or it can be parallel and promote the maintenace and the expansion of the monolayer [43]. The latter possibility makes the morphology of the epithelium more unstable, since there are more instability modes for a thin monolayer than for a stratified one. More precisely, the instability wavelength is given by the thickness of the monolayer [44, 45]. The orthogonal plane of division may be related to asymmetric cell division in which 2 different cells are produced: a proliferating basal cell with a stem identity, and a suprabasal cell. Misorientation of the division axis could contribute to cancer development by disorganising the tissue and provoking tissue metastasis [46]. Orientation of the mitotic spindle requires mechanical cues [47].

We write the evolution equation for *ϕ, c*, and the polarisation vector **p**, where *c* has the same meaning as in the previous sections. As before, the fluxes are represented by **v, v**_*c*_, *G*, Γ, *r*_*c*_, and we focus only on some specific couplings. Friction, viscosity, and growth are controlled by the parameters *ξ, η, g*_0_, and *γ* is the rotational viscosity. Active couplings between the polar source term, the proliferation term and the velocity are controlled by the parameters *ζ*_0_ and *ζ*_1_. The equations can be easily derived from Section 5 where the tensor **Q** has to be replaced by the vector **p**. We first explicit the free-energy, which, as in Section 5, is the sum of the Cahn-Hilliard Flory Huggins free energy, the polarity free energy, and the chemical free energy. *f* = *f*_*F H*_ + *f*_*P*_ + *f*_*c*_ where *f*_*F H*_ is written as above according to Eq. 8, and *f*_*c*_ as Eq. 13. Only *f*_*P*_ differs and reads:

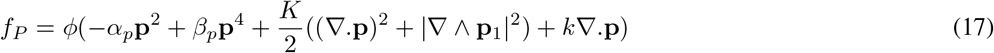

If the orientation of the cells is determined by the boundaries, i.e. cells spontaneously orient themselves parallel or perpendicular to boundaries, then *f*_*N*_ is coupled to *∇ϕ*. In this case the nematic free energy: *f*_*P*_ is modified and becomes:

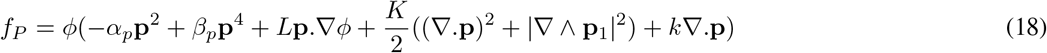

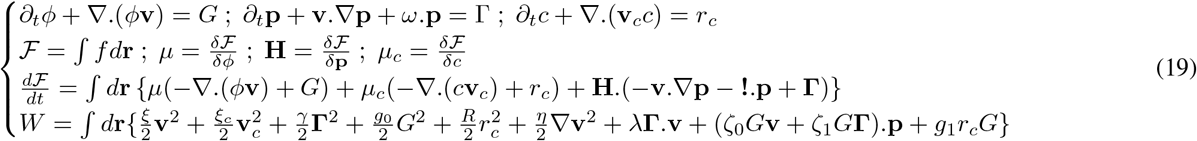

where *ω*_*αβ*_ = (*∂*_*α*_*v*_*β*_*− ∂*_*β*_*v*_*α*_)*/*2. We assume that activity is controlled by the molecules chemical potential, so that *g*_1_*/r*_*c*_ *→*0.

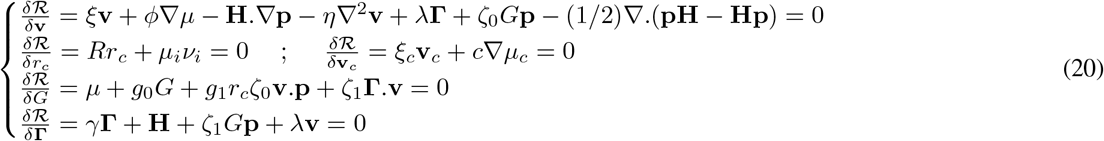

It comes:

If the plane of division is perpendicular to the cell polarity, the sign of *ζ*_0_ depends on whether the cell is extruded towards the basal side or the apical side. Conversely, if the plane of division is along the polarity, the cell divides along an axis perpendicular to its polarisation direction, so that the coupling *ζ*_0_*G***v.p** must be rewritten as a traceless tensor product *ζ*_0_*G*(**vv***−***I** Tr(**vv**)*/d*) : (**pp** *−***I** Tr(**pp**)*/d*) where the sign of *ζ*_0_ determines whether the orientations of velocity and tissue polarity are normal or aligned. In this case polar cells include features of merely nematic cells. In fact, active nematic stress has often been introduced as contributing to the stress in polar cells populations [48, 49].

In the following example, we show how coupling between growth and velocity can lead to instabilities of a tissue front when proliferation and cell death are related to nutrients.

### Polar front instability

We study a proliferating tissue in presence of a source of nutrients. This situation could be observed for instance in cancer where solid tumor proliferation and death depend notably on the availability of nutrients. The tissue is polar (*α*_*p*_ *>* 0), and its polarity is normal to its surface (*L >* 0). Cells have an orientation only when the tissue is dense enough: 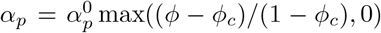. We start from a tissue with a plane interface between a dense and a dilute cellular phase (Fig. 4(a)). A source of nutrients is located on the top boundary of the simulation domain *y* = *L*, and the nutrients concentration is normalized by its value at *y* = *L* where *n*(*x, L*) = 1. The nutrients diffuse rapidly compared to the tissue evolution timescale, so that we can write the diffusion-consumption equation for the nutrient concentration as 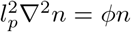 where *l*_*p*_ is the nutrient penetration length in the tissue. The nutrient concentration decreases from its source value in the dense phase (Fig. 4(b)). Cells proliferate or die depending on the nutrient concentration with a growth rate *G* = *ϕ*(*n−n*_0_)*/τ*_*G*_ (Fig. 4(c)). The presence of a polarity allows for a coupling between growth and velocity through the parameter *ζ*_0_. In absence of this coupling, no instability is observed on the interface (Fig. 4(d)). However, when *ζ*_0_ *<* 0, an instability of the proliferating front occurs.(Fig. 4(e)). This instability stems from the fact the velocity of the proliferating layer is in the direction of the polarity and in the direction of the dilute phase, whereas the cell death located deep in the tissue induces a velocity in the opposite direction.

**Figure 4.**
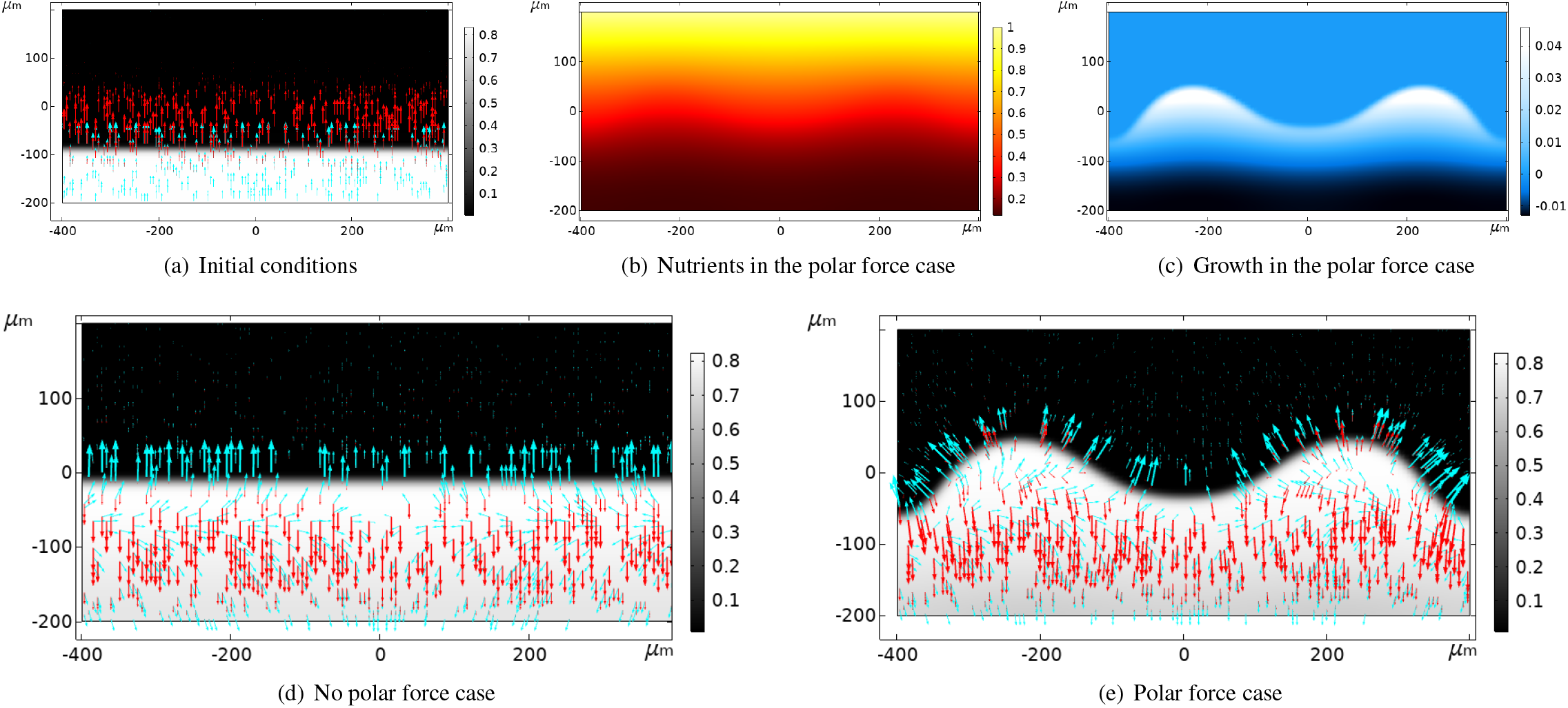
Comparison of a proliferating front in a polar force case and non polar force case. **4(a)**: initial condition for the numerical study. The polarity and velocity are indicated respectively with cyan and red arrows. The gray scale reflects the tissue mass fraction. **4(b)**: nutrients heatmap at *t* = 250 h for the polar case. **4(c)**: growth rate heatmap at *t* = 250 h for the polar case. **4(d)-4(e)**: mass fraction heatmap, polarity and velocity field for the apolar and polar cases at *t* = 250 h.

We now present a generalization of our formalism when different species interact in the mixing.

## 8 Formalism for different species

The previous derivation, limited to one tissue, can be extended to several species and tissues, provided that the mutual interactions between them are specified. We may have in mind the case of a tumour nest in its stroma but also the case of a growing epithelium growing between extracellular matrix called mesenchyme. Here is an example of such a modelling, in a growing tissue embedded in an active nematic shell: a classic case mimicking a desmo-plastic tumour in the pancreas or lung tumours. The belt appears due to the fibroblast population, resident in the microenvironment or as an activation of the immune inflammation. We show how activity in a nematic belt around proliferating cells potentially inhibits (in the case of contractile cells) or stimulates (in the case of extensile cells) e proliferating cells. In addition, we show how proliferative cells can generate active stresses related to the nematic component in their neighbourhood, and how these active stresses influence their own growth. Although we carry out this study in 2 dimensions, possibly corresponding to an experiment on a plate, this modelling captures a phenomenon that can occur in 3-dimensional spheroid both *in vitro* and *in vivo*.

The growing tissue is indexed by *g* and the active nematic tissue by *n*. The surrounding medium is indexed by *m*. So we have the mass fractions *ϕ*_*g*_, *ϕ*_*n*_, *ϕ*_*m*_ related by *ϕ*_*g*_ + *ϕ*_*n*_ + *ϕ*_*m*_ = 1. We only need to deal with the evolution equation for the growing and nematic components. The velocity of the growing tissue and of the nematic tissue are written as **v**_*g*_ and **v**_*n*_. The growth rate of the growing tissue is written with *G*. The symmetric and traceless nematic matrix of the nematic tissue is. **Q**_*αβ*_ and its source term writes **Γ**_*αβ*_. The evolution equations read:

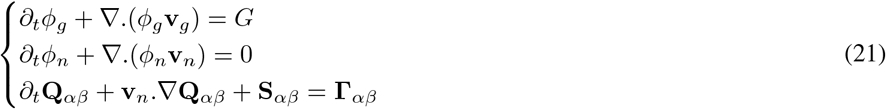

where **S**_*αβ*_ = **Q**_*αk*_**Ω**_*kβ*_*−* **Ω**_*αk*_**Q**_**kfi**_ and **Ω**_*αβ*_ = (*∂*_*β*_*v*_*nα*_ *− ∂*_*α*_*v*_*nβ*_)*/*2.

We now write the free-energy density of the mixture. It is a sum of a pure density term and a nematic contribution: *f* = *f*_*ϕ*_ + *f*_*N*_. The first part of the free energy is written as a Flory-Huggins/Cahn-Hilliard type of free-energy density. We assume that both tissues are cohesive with a negative square contribution but also attract each other, so that each forms a dense phase, which remains in contact with the other. It comes:

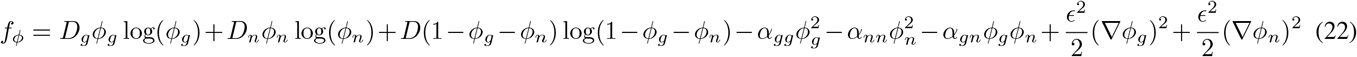

The nematic part of the free energy density includes a term that gives a natural orientation for the nematic cell type, and another term that prescribes this orientation with respect to the boundaries of this second cell type. It is written: as in the folllowing

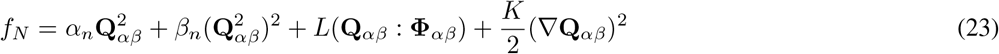

where *α*_*n*_ depends on the mass fraction of nematic cells: *α*_*n*_ = *α*_*n*0_(*ϕ− ϕ*_*c*_) and *β*_0_ *<* 0, so that the nematic cells acquire an average nematic order beyond the mass fraction *ϕ*_*c*_ for *ϕ > ϕ*_*c*_.

We write down the free energy and the chemical potentials and the time variation of the free energy resulting from this free energy density:

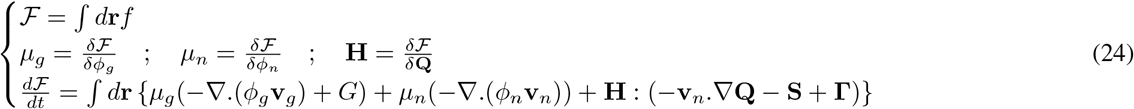

In our system, the fluxes are **v**_*g*_, **v**_*n*_, *G*, **Γ** as well as two advancement rates which are considered constant: the ATP activity advancement rate for the nematic cell type *r*_*n*_, and the growth related advancement rate *r*_*g*_ for the proliferating cell type. The dynamics are governed by a dissipation function which depends on the type of interactions present. In a first example, we show the case of proliferating cells in an active nematic belt, and in a second example, we show the case of active proliferating cells in a passive nematic belt.

### 8.1 Proliferating cells in an active nematic belt

The couplings include friction with the substrate, friction between the 2 cell types, as well as a coupling between the velocity and the nematic activity for the nematic cell type and coupling between growth and growth-related advancement rate for the proliferating cell type. It comes:

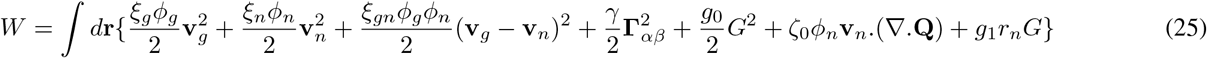

The leading equations for the fluxes are obtained after the minimisation of the Rayleighian with respect to these fluxes: *ℛ* = 𝒲 + *d ℱ /dt*. We obtain:

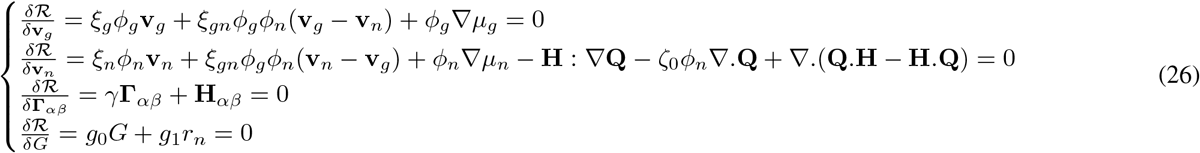

Without loss of generality, we choose a modified logistic expression which implies the growth rate: *G* = (*ϕ*^*max*^*− ϕ*_*g*_ *−ϕ*_*n*_) max(*ϕ*_*g*_*−ϕ*^*min*^, 0)*/τ*_*G*_ that implies that growth saturates and apoptosis occurs for *ϕ*_*g*_ *> ϕ*^*max*^ and that cells are quiescent for *ϕ*_*g*_ below *ϕ*_*min*_.

We now show the different scenarios that are observed in Fig. 5. In Fig. 5(a) the tissue has a structure with the proliferative cells (in red) embedded in a nematic component (in green, with the lines following the nematic orientation). The extent of growth depends on the type of stress in the nematic belt (Fig. 5(b)), as we implement a logistic growth rate. Contractile stress compresses the proliferating cells close to the threshold for cell death *ϕ*^*max*^ (Fig. 5(c)). Conversely, tensile stress tends to reduce the mass fraction for the proliferating cells as their tissue portion is under tension. In all the cases a nematic order surrounds the proliferating cells (Fig. 5(d)).

**Figure 5.**
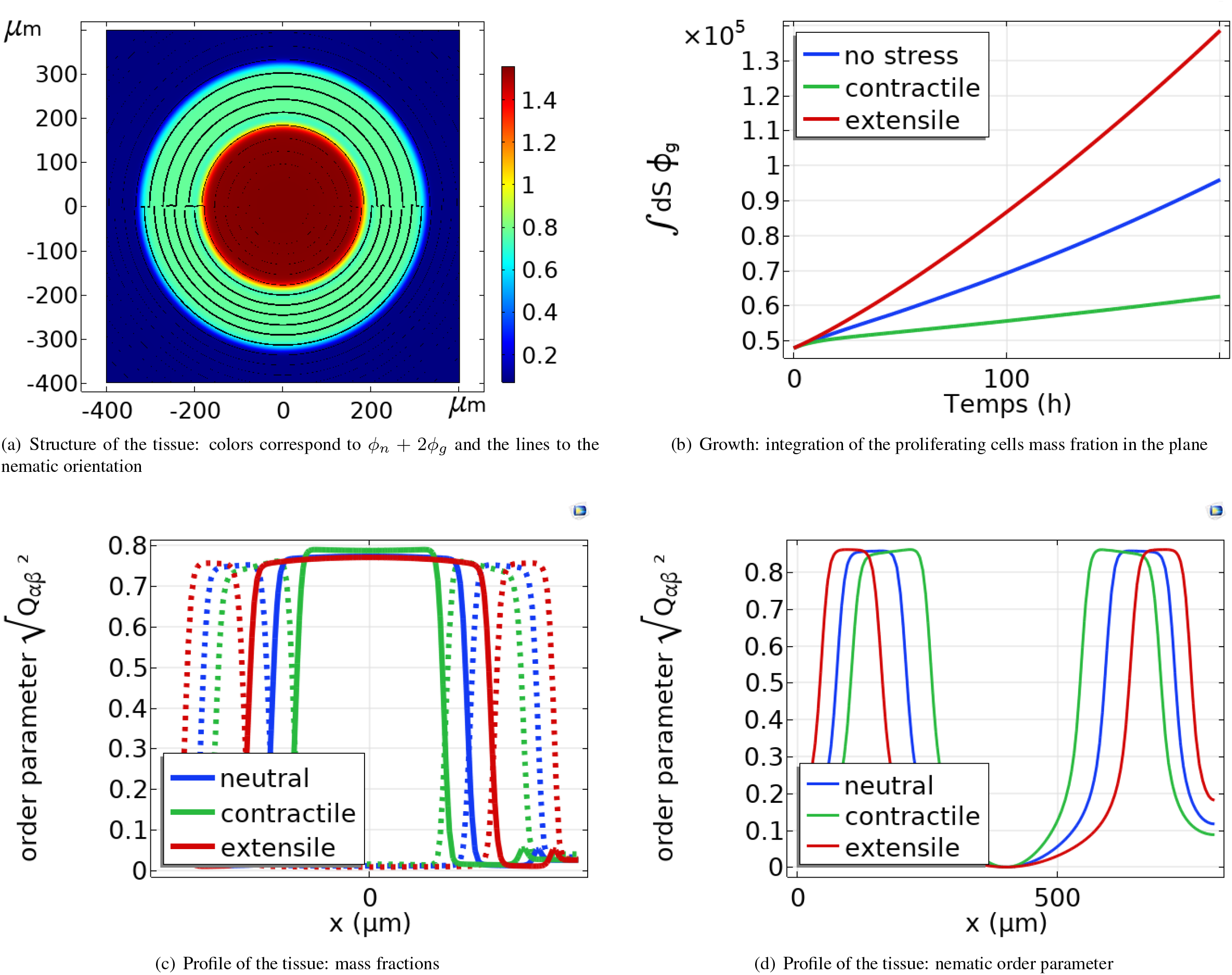
Proliferating cells sourrounded by active nematic cells. 5(a): Structure of the tissue: colors correspond to *ϕ*_*n*_ + 2*ϕ*_*g*_ and allow to identify the different tissue components. The lines correspond to the nematic orientation **n** = (cos(atan2(*Q*_*xy*_, *Q*_*xx*_)*/*2), sin(atan2(*Q*_*xy*_, *Q*_*xx*_)*/*2)). 5(b): Integration of the mass fraction over the system in function of time in *m*^2^. 5(c): Cut of the numerical study window showing the profile of mass fraction at *t* = 200 h for each scenario of extensile, contractile or no active stress. 5(d): Cut of the numerical study window showing the profile of the nematic order parameter at *t* = 200 h for each scenario of extensile, contractile or no active stress.

### 8.2 Active proliferating cells in a nematic belt

In this second example, we introduce active stress in the proliferating cells and not in the nematic cells. The proliferating cells apply stress to themselves in the presence of a nematic environment. The dissipation function becomes:

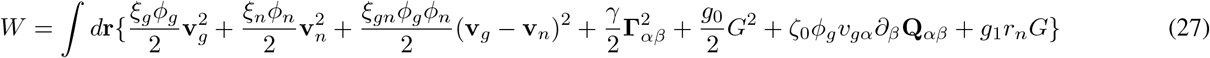

and the equations relating the fluxes read:

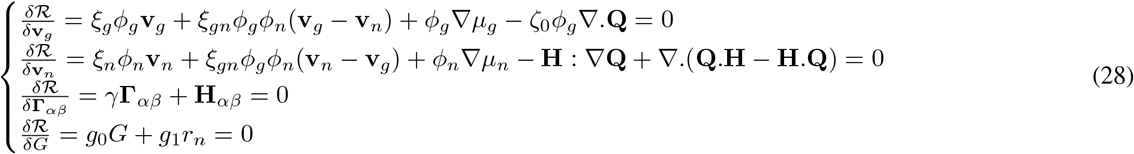

As before, we choose the modified logistic growth rate *G* = (*ϕ*^*max*^*− ϕ*_*g*_*− ϕ*_*n*_) max(*ϕ*_*g*_*− ϕ*^*min*^, 0)*/τ*_*G*_. Active stresses still lead to an increase in total growth in the case of a tensile stress, and to a decrease in the case of a contractile stress (Fig. 6(a)). These variations are related to the mass fraction of proliferating cells in the bulk of their dense phase (Fig. 6(b)).

**Figure 6.**
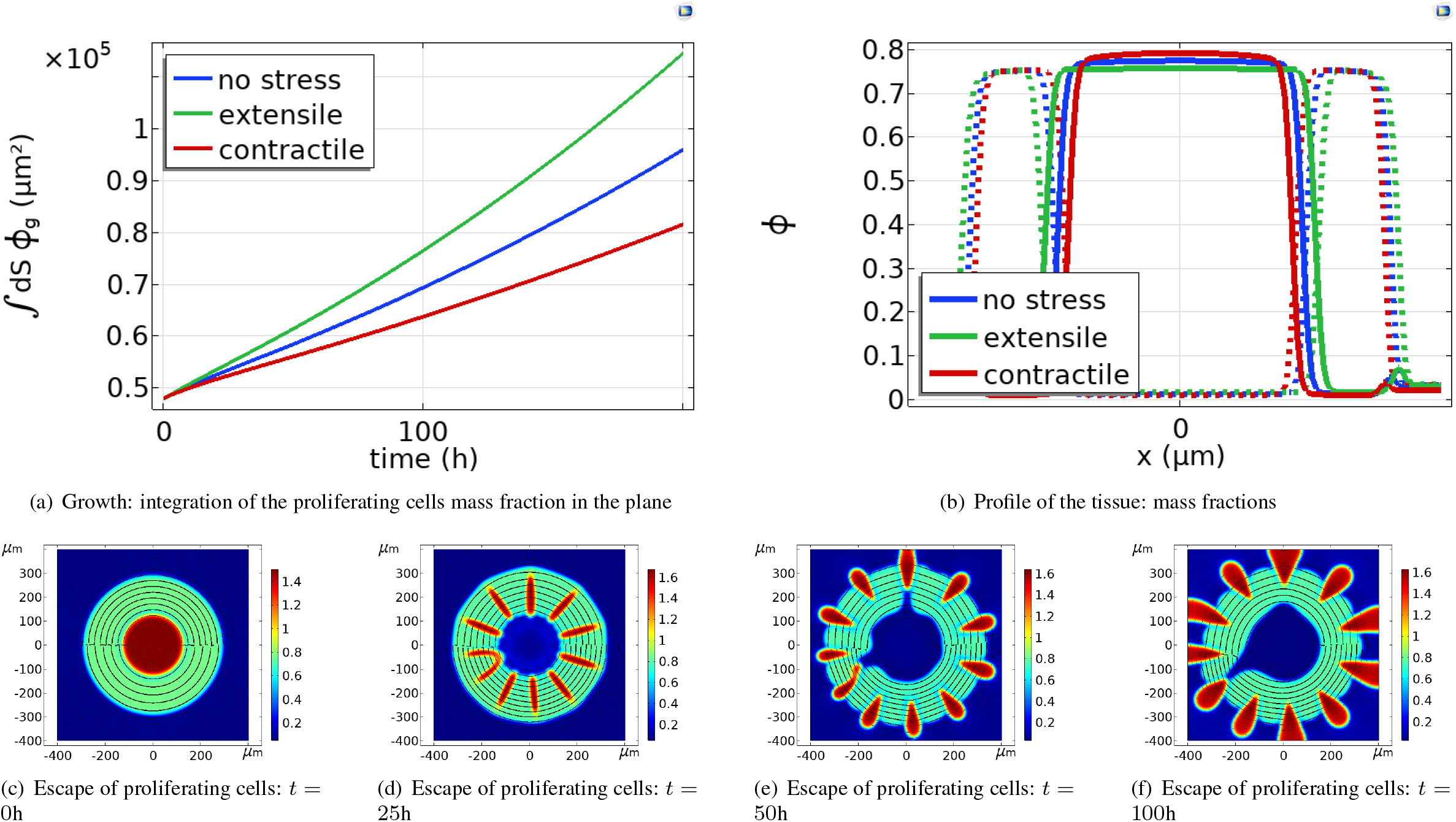
Active proliferating cells sourrounded by passive nematic cells. 6(a): Integration of the mass fraction over the system in function of time in *m*^2^. 6(b): Cut of the numerical study window showing the profile of the mass fraction at *t* = 200 h for each scenario of tensile, contractile or no active stress. 6(c)-6(f): Escape of proliferating cells in presence of a tensile active stress. The color legend corresponds to *ϕ*_*n*_ + 2*ϕ*_*g*_. Thus, proliferating cells are in red and nematic cells are in green.

Further increases in tensile stress leads to the escape of the proliferating cells from the ring of nematic cells (Fig. 6(c)-6(f)).

## 9 Conclusion

In this article, we have shown how to easily derive the different possible physical couplings in a given complex system involving different partners as cells, tissues, chemicals, fluids in the presence of growth by Onsager’s Variational approach and the definition of a Rayleighian [21, 20]. Onsager’s Variational Principle is equivalent to the expansion of fluxes in terms of forces, that has been used to derive the generalized hydrodynamics of similar problems [41, 6]. However, in the variational principle, fluxes are more clearly defined, and the force balance equations are easily derived. We have applied this method to different cases of proliferating systems in two dimensions, with a special focus on cancer cells. However, this method can be generalised to growing tissues in general as growing epithelia during morphogenesis and embryogenesis and to other pathologies as fibrosis. We showed how to systematically derive the equations for apolar, polar, and nematic tissues in presence or not of activity. We showed how growth can depend on those properties. Notably, we showed how instabilities can be triggered by activity or more generally by introducing chemicals in the system. Although we only applied our formalism to 2D simulations, a wide panel of scenarios are revealed. For instance, we showed how nematic cells can grow in a defined direction or on the contrary present instabilities depending on their specific energy and activity couplings. Similarly, we showed how despite the surface tension of the tissue, the presence of nutrients and the balance between cell proliferation and cell death destabilizes the interface between the dense and dilute phases. The addition of multiple cell species in the tissue allows for a large numbers of coupling. Remarkably, non polar/nematic species can use the nematic properties of other cell species in order to acquire an activity. Indeed, it was shown in the section on apolar/anematic tissues that although there can exist a coupling between the growth rate and activity, coupling between velocity and activity requires a directionality in the tissue that can be proposed only by gradients of cell density in absence of nematic or polar properties.

**Table Sup. 1:** Parameter values in the different numerical studies. *β* = 0 in all the numerical studies. *k* = 0 in the polar numerical study.

| Figure<br>Parameters | 3(a) | 3(b) | 3(c) | 3(d) | 3(e) | 3(f) | 4 | 5 | 6(a) | 6(c) |
| --- | --- | --- | --- | --- | --- | --- | --- | --- | --- | --- |
| $D_0, D$ ($kg.\mu m^{-1}.h^{-2}$) | $10^4$ | $10^4$ | $10^4$ | $10^4$ | $10^4$ | $10^4$ | $10^4$ | $10^4$ | $10^4$ | $10^4$ |
| $D_1, D_g$ ($kg.\mu m^{-1}.h^{-2}$) | $2 \times 10^3$ | $2 \times 10^3$ | $2 \times 10^3$ | $2 \times 10^3$ | $2 \times 10^3$ | $3 \times 10^3$ | $2 \times 10^3$ | $2 \times 10^3$ | $2 \times 10^3$ | $2 \times 10^3$ |
| $D_n$ ($kg.\mu m^{-1}.h^{-2}$) | X | X | X | X | X | X | X | $2 \times 10^3$ | $2 \times 10^3$ | $2 \times 10^3$ |
| $\alpha, \alpha_{gg}$ ($kg.\mu m^{-1}.h^{-2}$) | $1.5 \times 10^4$ | $1.5 \times 10^4$ | $1.5 \times 10^4$ | $1.5 \times 10^4$ | $1.5 \times 10^4$ | $1.5 \times 10^4$ | $1.6 \times 10^4$ | $1.4 \times 10^4$ | $1.4 \times 10^4$ | $1.4 \times 10^4$ |
| $\alpha_{gn}$ ($kg.\mu m^{-1}.h^{-2}$) | X | X | X | X | X | X | X | $1.7 \times 10^4$ | $1.7 \times 10^4$ | $1.7 \times 10^4$ |
| $\alpha_{nn}$ ($kg.\mu m^{-1}.h^{-2}$) | X | X | X | X | X | X | X | $1.4 \times 10^4$ | $1.4 \times 10^4$ | $1.4 \times 10^4$ |
| $\epsilon^2$ ($kg.\mu m.h^{-2}$) | $2.2 \times 10^6$ | $2.2 \times 10^6$ | $2.2 \times 10^6$ | $2.2 \times 10^6$ | $2.2 \times 10^5$ | $2.2 \times 10^6$ | $3.6 \times 10^5$ | $3.6 \times 10^5$ | $3.6 \times 10^5$ | $3.6 \times 10^5$ |
| $\xi, \xi_g$ ($kg.\mu m^{-3}.h^{-1}$) | 0 | 0 | 0 | 0 | 0 | 1 | 10 | 1 | 1 | 1 |
| $\xi_n$ ($kg.\mu m^{-3}.h^{-1}$) | X | X | X | X | X | X | X | 1 | 1 | 1 |
| $\xi_{gn}$ ($kg.\mu m^{-3}.h^{-1}$) | X | X | X | X | X | X | X | 10 | 10 | 10 |
| $\eta$ ($kg.\mu m^{-1}.h^{-1}$) | $10^3$ | $10^3$ | $10^3$ | $10^3$ | $10^5$ | $10^3$ | 0 | 0 | 0 | 0 |
| $\zeta_0$ ($kg.\mu m^{-1}.h^{-1}$) | 0 | 0 | 0 | $-10^3$ | $0, \pm 10^6$ | -1 | $0, 10^2$ | $0, \pm 10^3$ | $0, \pm 10^3$ | $-10^4$ |
| $\zeta_1$ ($kg.\mu m^{-1}.h^{-1}$) | 0 | 0 | -0.1 | -0.2 | 0 | 0 | 0 | 0 | 0 | 0 |
| $\lambda, \lambda_0$ ($kg.\mu m^{-1}.h^{-1}$) | $7.8 \times 10^{-1}$ | $-7.8 \times 10^{-1}$ | 0 | 0 | 0 | 0 | 0 | 0 | 0 | 0 |
| $\gamma$ ($kg.\mu m^{-1}.h^{-1}$) | $10^{-2}$ | $10^{-2}$ | $10^{-2}$ | $10^{-2}$ | $10^{-2}$ | $10^{-4}$ | $10^{-4}$ | $10^{-2}$ | $10^{-2}$ | $10^{-2}$ |
| $\tau_G$ (h) | 10 | 10 | 10 | 10 | 30 | 20 | 4 | 9 | 9 | 9 |
| $\alpha_n, \alpha_{n0}, \alpha_p^0$ ($kg.\mu m^{-1}.h^{-1}$) | 0 | 0 | 0 | 0 | 0 | $10^{-2}$ | $5 \times 10^{-3}$ | $10^{-1}$ | $10^{-1}$ | $10^{-1}$ |
| $\beta_n, \beta_p$ ($kg.\mu m^{-1}.h^{-1}$) | $2 \times 10^{-2}$ | $2 \times 10^{-2}$ | $2 \times 10^{-2}$ | $2 \times 10^{-2}$ | $2 \times 10^{-2}$ | $2 \times 10^{-2}$ | $2.5 \times 10^{-3}$ | $2 \times 10^{-1}$ | $2 \times 10^{-1}$ | $2 \times 10^{-1}$ |
| $\epsilon_Q^2, K$ ($kg.\mu m.h^{-1}$) | 9 | 9 | 9 | 9 | 9 | 9 | 1 | 3.6 | 3.6 | 3.6 |
| $L$ ($kg.\mu m.h^{-1}$) | 0 | 0 | 0 | 0 | $5 \times 10^{-2}$ | 0 | 0.5 | $10^{-1}$ | $10^{-1}$ | $10^{-1}$ |

This work has focused on 2D applications, and it would be interesting to continue this study in 3D structures. For this purpose, nematic tissues on 2D curved surfaces represent an interesting intermediate state, whose active and passive behaviour has been studied in the case of shells and vesicles, and the large variety of scenarios has not yet been fully explored [50, 51, 52, 53, 54, 55, 56]. In addition, fluctuations have been shown to be able to induce nematic order and stresses, so their role in nematic-related growth remains to be elucidated [57]. Last, an interesting question is the continuous modelling of irregular cell shapes with no particular orientation, that controls rigidity transition in vertex models, notably in the context of EMT [58].

In conclusion, the formalism that we have developed for tissues can be extended and applied to many biophysical processes occurring in healthy or pathological tissues.

## 10 Appendix: Parameters used in the simulations

The values of the parameters used in the different simulations are shown in Table.Sup. 1. These parameters are highly variable depending on the type of tissue studied. In our study, the values of the parameters related to the Cahn-Hilliard Flory-Huggins (CH-FH) free energy are found in the literature, as well as the values of the viscosity and the friction to the substrate in the cases we introduce them [59, 60, 1]. The other parameters are chosen to obtain the displayed patterns. Interestingly, the nematic and polar free-energy scales are found to be much lower than the CH-FH free-energy.

## 11 Author Contributions

The authors contribute equally.

## 12 Acknowledgements

J.A and M.BA acknowledges the financial support from ITMO Cancer of Aviesan within the framework of the 2021-2030 Cancer Control Strategy, on funds administrated by Inserm (PCSI 2021,MCMP 2022). J.A acknowledges the financial support from ANR COLLAMOEBOID. MBA would like to thank the Isaac Newton Institute for Mathematical Sciences for support and hospitality during the program “Uncertainty Quantification and stochastic modelling of materials”. This work was supported by: EPSRC Grant Number EP/R014604/1.

